# Geochemical and metagenomic characterization of Jinata Onsen, a Proterozoic-analog hot spring, reveals novel microbial diversity including iron-tolerant phototrophs and thermophilic lithotrophs

**DOI:** 10.1101/428698

**Authors:** Lewis M. Ward, Airi Idei, Mayuko Nakagawa, Yuichiro Ueno, Woodward W. Fischer, Shawn E. McGlynn

## Abstract

Hydrothermal systems, including terrestrial hot springs, contain diverse geochemical conditions that vary over short spatial scales due to progressive interaction between the reducing hydrothermal fluids, the oxygenated atmosphere, and in some cases seawater. At Jinata Onsen, on Shikinejima Island, Japan, an intertidal, anoxic, iron-rich hot spring mixes with the oxygenated atmosphere and seawater over short spatial scales, creating a diversity of chemical potentials and redox pairs over a distance ∼10 m. We characterized the geochemical conditions along the outflow of Jinata Onsen as well as the microbial communities present in biofilms, mats, and mineral crusts along its traverse via 16S rRNA amplicon and genome-resolved shotgun metagenomic sequencing. The microbial community changed significantly downstream as temperatures and dissolved iron concentrations decreased and dissolved oxygen increased. Near the spring source, visible biomass is limited relative to downstream, and primary productivity may be fueled by oxidation of ferrous iron and molecular hydrogen by members of the Zetaproteobacteria and Aquificae. Downstream, the microbial community is dominated by oxygenic Cyanobacteria. Cyanobacteria are abundant and active even at ferrous iron concentrations of ∼150 μM, which challenges the idea that iron toxicity limited cyanobacterial expansion in Precambrian oceans. Several novel lineages of Bacteria are also present at Jinata Onsen, including previously uncharacterized members of the Chloroflexi and Caldithrichaeota phyla, positioning Jinata Onsen as a valuable site for future characterization of these clades.

## Introduction

A major theme of environmental microbiology has been the enumeration of microbial groups that are capable of exploiting diverse chemical potentials (i.e. chemical disequilibria) that occur in nature (e.g. 8, 27, 115). Hot springs, with their varied chemical compositions, provide reservoirs of novel microbial diversity, where environmental and geochemical conditions select for lineages and metabolisms distinct from other Earth-surface environments (e.g. 4, 10, 28, 130, 131). In addition to their value as sources of microbial diversity, hot springs also provide valuable test beds for understanding microbial community processes driven by different suites of metabolisms (e.g. 52)—this in turn allows these systems to serve as process analogs and to provide a window into biosphere function during early times in Earth history, for example when the O_2_ content of surface waters was low or non-existent. In contrast to most surface ecosystems which are fueled almost entirely by oxygenic photosynthesis by plants, algae, and Cyanobacteria, hot spring microbial communities are commonly supported by lithotrophic or anoxygenic phototrophic organisms that derive energy and electrons for carbon fixation by oxidizing geologically sourced electron donors such as Fe^2+^, sulfide, arsenite, and molecular hydrogen (e.g. 36, 65, 71, 109, 130). These communities may therefore provide insight into the function of microbial communities on the early Earth or other planets, in which oxygenic photosynthesis may be absent or less significant and anoxygenic photosynthetic or lithotrophic metabolisms may play a larger role, resulting in overall lower rates of primary productivity (e.g. 14, 66, 101, 135, 136, 137).

Here, we present a geomicrobiological characterization of a novel Precambrian Earth process analog site: Jinata Onsen, on Shikinejima Island, Tokyo Prefecture, Japan. While a small number of metagenome-assembled genomes have previously been recovered from Jinata (129, 131), we describe here the first overall characterization of the geochemistry and microbial community of this site. This site supports sharp gradients in geochemistry that in some ways recapitulate spatially environmental transitions which occurred temporally during Proterozoic time. The modern, sulfate-rich, well-oxygenated ocean that we see today is a relatively recent state, typical only of only the last few hundred million years (e.g. 79). For the first half of Earth history, until ∼2.3 billion years ago (Ga), the atmosphere and oceans were anoxic (54), and the oceans were largely rich in dissolved iron but poor in sulfur (124). Following the Great Oxygenation Event ∼2.3 Ga, the atmosphere and surface ocean accumulated some oxygen, and the ocean transitioned into a stratified state with oxygenated surface waters and anoxic deeper waters, rich in either dissolved iron or sulfide (92). At Jinata Onsen, this range of geochemical conditions is recapitulated over just a few meters, providing an ideal space-for-time analog to test hypotheses of how microbial diversity and productivity may have varied as environmental conditions changed through Earth history.

At Jinata hot spring, anoxic, iron-rich hydrothermal fluids feed a subaerial spring that flows into a small bay, and mixes with seawater over the course of a few meters. Over its course the waters transition from low-oxygen, iron-rich conditions analogous to some aspects of the early Proterozoic oceans, toward iron-poor and oxygen-rich conditions typical of modern coastal oceans. In upstream regions of the stream where oxygenic Cyanobacteria are at very low abundance, biomass is visibly sparse; however, downstream, biomass accumulates in the form of thick microbial mats containing abundant Cyanobacteria. Visible differences in accumulation and appearance of biomass across the temperature and redox gradient establish the hypothesis that microbial community composition, as well as the magnitude and metabolic drivers of primary productivity, varies along the spring flow. To begin testing this hypothesis and to provide a baseline description of the geochemistry and microbiology of this site in support of future investigation, we performed geochemical measurements, 16S rRNA amplicon sequencing, and genome-resolved metagenomic sequencing to recover draft genomes of diverse novel microbial lineages that inhabit Jinata Onsen.

## Materials and Methods

### Geological context and sedimentology of Jinata

Jinata Onsen is located at 34.318 N, 139.216 E on the island of Shikinejima, Tokyo Prefecture, Japan. Shikinejima is part of the Izu Islands, a chain of volcanic islands that formed in the last few million years along the northern edge of the Izu-Bonin-Mariana Arc (58). Shikinejima is formed of Late Paleopleistocene-to-Holocene non-alkaline felsic volcanics and Late-Miocene to Pleistocene non-alkaline pyroclastic volcanic flows, with Jinata Onsen located on a small bay on the southern side of the island (Figure 1).

**Figure 1:**
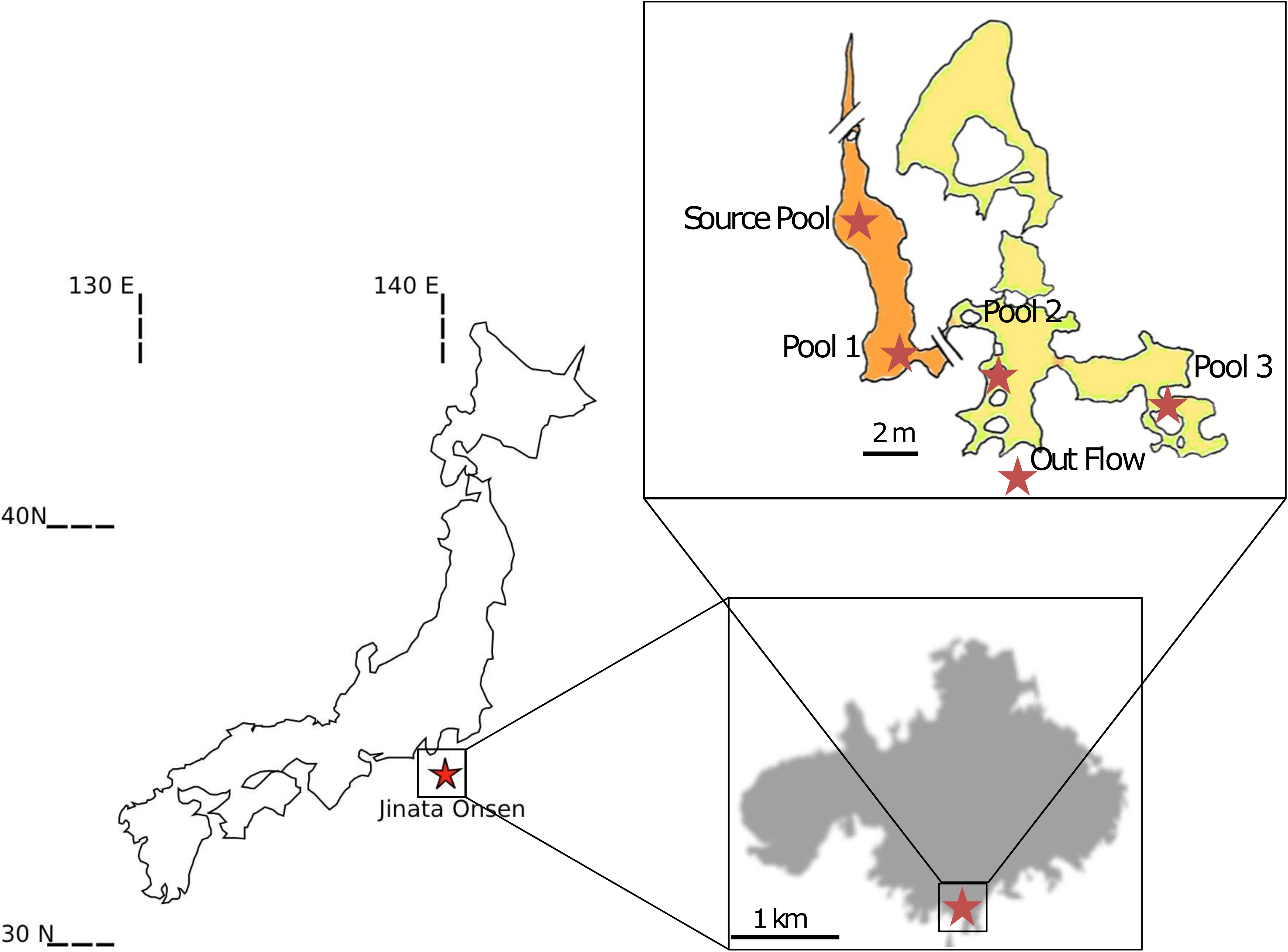
Location of Jinata Onsen on Shikinejima Island, Japan, and inset overview sketch of field site with sampling localities marked.

### Sample collections

Five sites were sampled at Jinata Onsen: the Source Pool, Pool 1, Pool 2, Pool 3, and the Outflow (Figure 1, Figure 2). During the first sampling trip in January 2016, two whole community DNA samples were collected from each site for 16S rRNA amplicon sequencing. During the second sampling trip, additional DNA was collected from the Source Pool and Pool 2 for shotgun metagenomic sequencing along with gas samples for qualitative analysis. Samples for quantitative gas analysis were collected in October 2017 and April 2018.

**Figure 2:**
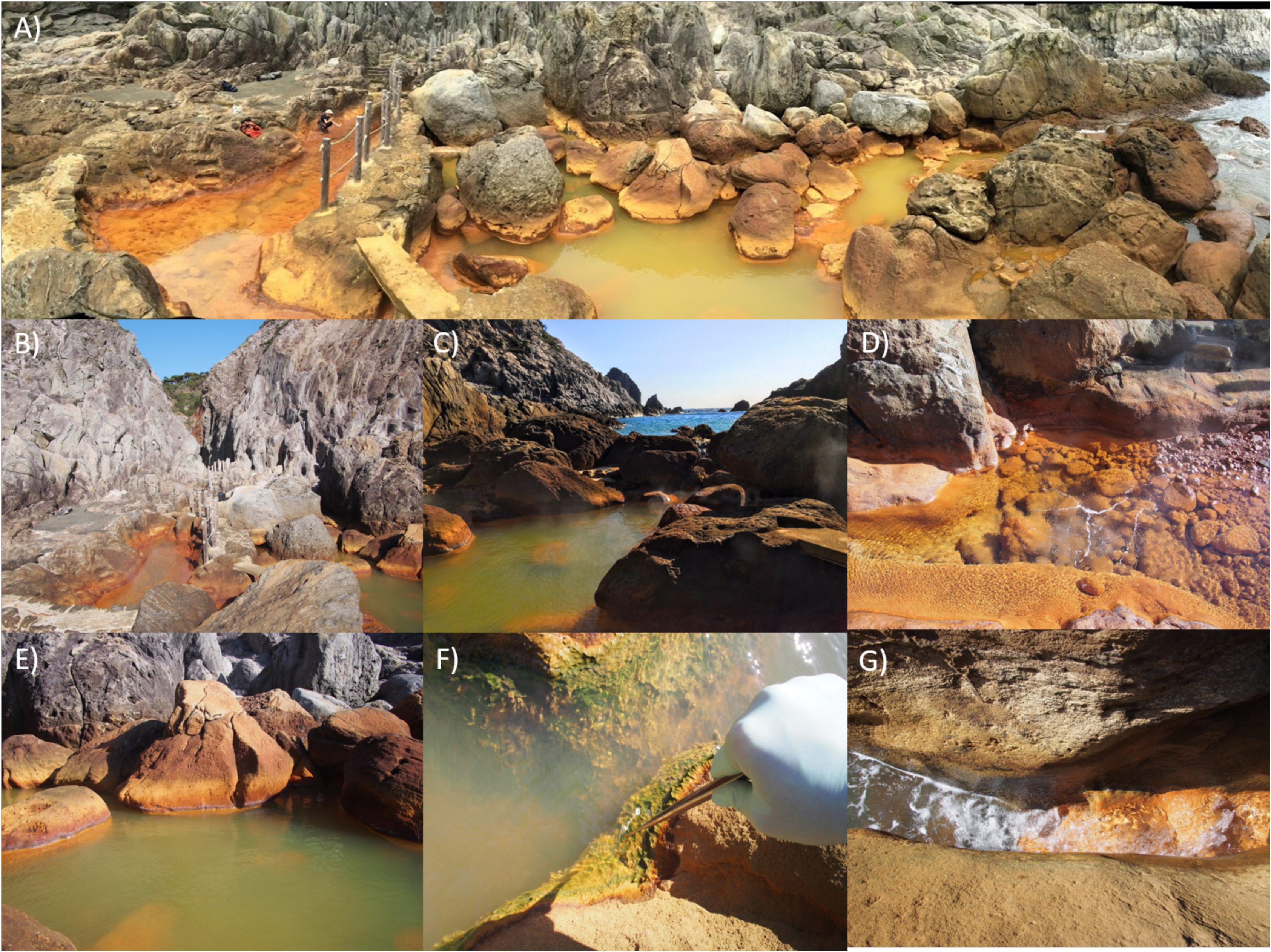
Representative photos of Jinata. A) Panorama of field site, with Source Pool on the left (Pool 1 below), Pool 2 and 3 in the center, and Out Flow to the bay on the right. B) Undistorted view north up the canyon. C) Undistorted view south toward the bay, overlooking Pool 2. D) Source Pool, coated in floc-y iron oxides and bubbling with gas mixture containing CO_2_, CH_4_ and trace, potentially variable, H_2_. E) Pool 2, with mixture of red iron oxides and green from Cyanobacteria-rich microbial mats. F) Close up of textured microbial mats in Pool 3. G) Close up of Out Flow, where hot spring water mixes with ocean water. Reprinted with permission from (131).

Samples were collected as mineral scrapings of loosely attached, fluffy iron oxide coating from surfaces and clasts upstream (Source Pool and Pool 1) and as samples of microbial mat downstream (Pools 2 and 3, and Outflow) using sterile forceps and spatulas (∼0.25 cm^3^ of material). Immediately after sampling, cells were lysed and DNA preserved with a Zymo Terralyzer BashingBead Matrix and Xpedition Lysis Buffer. Lysis was achieved by attaching tubes to the blade of a cordless reciprocating saw (Black & Decker, Towson, MD) and operating for 1 minute. Aqueous geochemistry samples consisted of water collected with sterile syringes and filtered through a 0.2 μm filter. Gas samples were collected near sites of ebullition emerging from the bottom of the Source Pool; collection was done into serum vials by water substitution, and then sealed underwater to prevent contamination by air.

### Geochemical analysis

Dissolved oxygen (DO), pH, and temperature measurements were performed *in situ* using an Extech DO700 8-in-1 Portable Dissolved Oxygen Meter (FLIR Commercial Systems, Inc., Nashua, NH). Iron concentrations were measured using the ferrozine assay (114) following acidification with 40 mM sulfamic acid to inhibit iron oxidation by O_2_ or oxidized nitrogen species (69). Ammonia/ammonium concentrations were measured using a TetraTest NH_3_/NH_4_ ^+^ Kit (TetraPond, Blacksburg, VA) following manufacturer’s instructions but with colorimetry of samples and NH_4_Cl standards quantified with a Thermo Scientific Nanodrop 2000c spectrophotometer (Thermo Fisher Scientific, Waltham, MA) at 700 nm to improve sensitivity and accuracy. Anion concentrations were measured via ion chromatography on a Shimadzu Ion Chromatograph (Shimadzu Corp., Kyoto, JP) equipped with a Shodex SI-90 4E anion column (Showa Denko, Tokyo, JP).

Presence of H_2_ and CH_4_ in gas samples was initially qualitatively determined by comparison to standards with a Shimadzu GC-14A gas chromatograph within 12 hours of collection to minimize oxidation of reduced gases. Subsequent gas samples were analyzed following methods from (116). In brief, samples were analyzed using a gas chromatograph (GC-4000, GL Sciences) equipped with both a pulsed discharge detector (PDD) and a thermal conductivity detector (TCD). The GC was equipped with a ShinCarbon ST packed column (2 m × 2.2 mm ID, 50/80 mesh) connected to a HayeSepo Q packed column (2 m × 2.2 mm ID, 60/80 mesh) to separate O_2_, N_2_, CO_2_, and light hydrocarbons. Temperature was held at 40°C for 6 minutes before ramping up to 200°C at 20°C/min. This temperature was held for 6 minutes before ramping up to 250°C at 50°C/min before a final hold for 15 minutes. The value of standard errors (SE) were determined by replicate measurement of samples. The detection limit was on the order of 1nmol/cc for H_2_ and CH_4_.

Water samples for dissolved inorganic carbon (DIC) and dissolved organic carbon (DOC) concentration measurements were collected with sterile syringes and transferred after filtering through a 0.2 μm filter to pre-vacuumed 30 mL serum vials which were sealed with butyl rubber septa and aluminum crimps.

DIC and DOC concentrations in water samples were analyzed by measuring CO_2_ in the headspace of the sampled vials after the reaction of sample with either phosphoric acid for DIC or potassium persulfate for DOC with a Shimadzu GC-14A gas chromatograph. Sodium bicarbonate standards and glucose standards were used for making calibration curves to quantify DIC and DOC concentrations, respectively.

### 16S rRNA and metagenomic sequencing and analysis

Sequencing and analysis of 16S rRNA amplicon data followed methods from (130). Following return to the lab, bulk environmental DNA was extracted and purified with a Zymo Soil/Fecal DNA extraction kit. The V4-V5 region of the 16S rRNA gene was PCR amplified using archaeal and bacterial primers 515F (GTGCCAGCMGCCGCGGTAA) and 926R (CCGYCAATTYMTTTRAGTTT) (15). DNA was quantified with a Qubit 3.0 fluorimeter (Life Technologies, Carlsbad, CA) following the manufacturer’s instructions following DNA extraction and PCR steps. Successful amplification of all samples was verified by viewing on a gel after initial pre-barcoding PCR (30 cycles). Duplicate PCR reactions were pooled and reconditioned for five cycles with barcoded primers. Samples for sequencing were submitted to Laragen (Culver City, CA) for analysis on an Illumnia MiSeq platform. Sequence data were processed using QIIME version 1.8.0 (15). Raw sequence pairs were joined and quality-trimmed using the default parameters in QIIME. Sequences were clustered into de novo operational taxonomic units (OTUs) with 99% similarity using UCLUST open reference clustering protocol (26). Then, the most abundant sequence was chosen as representative for each *de novo* OTU (125). Taxonomic identification for each representative sequence was assigned using the Silva-132 database (94) clustered separately at 99% and at 97% similarity. Singletons and contaminants (OTUs appearing in the negative control datasets) were removed. 16S rRNA sequences were aligned using MAFFT (62) and a phylogeny constructed using FastTree (93). Alpha diversity was estimated using the Shannon Index (100) and Inverse Simpson metric (1/D) (48, 104). Assessment of sampling depth was estimated using Good’s Coverage (37). All statistics were calculated using scripts in QIIME and are reported at the 99% and 97% OTU similarity levels. Multidimensional scaling (MDS) analyses and plots to evaluate the similarity between different samples and environments were produced in R using the vegan and ggplot2 packages (86, 95, 139).

Following initial characterization via 16S rRNA sequencing, four samples were selected for shotgun metagenomic sequencing: JP1-A and JP3-A from the first sampling trip, and JP1L-1 and JP2-1 from the second sampling trip. Purified DNA was submitted to SeqMatic LLC (Fremont, CA) for library preparation and 2×100bp paired-end sequencing via Illumina HiSeq 4000 technology. Samples JP1-A and JP3-A shared a single lane with two samples from another project, while JP1L-1 and JP2-1 shared a lane with one sample from another project.

Raw sequence reads from all four samples were co-assembled with MegaHit v. 1.02 (77) and genome bins constructed based on nucleotide composition and differential coverage using MetaBAT (59), MaxBin (140), and CONCOCT (2) prior to dereplication and refinement with DAS Tool (103) to produce the final bin set. Genome bins were assessed for completeness, contamination, and strain-level heterogeneity using CheckM (89), tRNA sequences found with Aragorn (73), and presence of metabolic pathways of interest predicted with MetaPOAP (134). Coverage was extracted using bbmap (11) and samtools (76). Genes of interest (e.g. coding for ribosomal, photosynthesis, iron oxidation, and electron transport proteins) were identified from assembled metagenomic data locally with BLAST+ (12 and were screened against outlier (e.g. likely contaminant) contigs as determined by CheckM using tetranucleotide, GC, and coding density content. Translated protein sequences of genes of interest were aligned with MUSCLE (25), and alignments manually curated in Jalview (138). Phylogenetic trees were calculated using RAxML (110) on the Cipres science gateway (83). Node support for phylogenies was calculated with transfer bootstraps by BOOSTER (74). Trees were visualized with the Interactive Tree of Life viewer (Letunic and Bork 2016). Because sequencing depth of each sample in the full metagenome was uneven, relative abundance of genes of interest between metagenomic datasets was normalized to the coverage of *rpoB* genes in each raw dataset as mapped onto the coassembly. Like the 16S rRNA gene, *rpoB* is a highly conserved, vertically-inherited gene useful for taxonomic identification of organisms but has the added advantage that it is only known to occur as a single copy per genome (17) and is more readily assembled in metagenomic datasets (e.g. 131). Presence and classification of hydrogenase genes was determined with HydDB (107). Taxonomic assignment of MAGs was made based on placement in a reference phylogeny built with concatenated ribosomal protein sequences following Hug et al. (50) and confirmed using GTDB-Tk (90). Optimal growth temperatures of MAGs was predicted based on proteome-wide 2-mer amino acid composition following methods from (78).

## Results

### Site description

The source water of Jinata Onsen emerges with low dissolved oxygen concentrations near our limit of detection, is iron-rich, and gently bubbles gas from the spring source (Table 1, Figure 1, Figure 2). Temperatures at the source are ∼63°C. Water emerges into the Source Pool, which has no visible microbial mats or biofilms (Figure 2D). Surfaces are instead coated with a fluffy red precipitate, likely a poorly ordered or short range-ordered ferric iron oxide phase such as ferrihydrite. Flow from the source is—at least in part—tidally charged, with the highest water levels and flow rates occurring at high tide. At low tide, flow rates drop and the water level of the Source Pool can drop by decimeters and portions of the Source Pool can drain during spring low tides. Downstream, the spring water collects into a series of pools (Pool 1-3) (Figure 2C,E-F), which cool sequentially (Figure 3, Supplemental Table 1). Pool 1 contains iron oxides like the Source Pool, but also develops macroscopic microbial streamers that are coated in iron oxides and thin veil-like layers of microorganisms overlaying iron oxide sediments—structures similar to those typically made by marine iron-oxidizing Zetaproteobacteria (e.g. 35). Streamers are very fine (mm-scale) and delicate (break apart on contact with forceps) but can reach several centimeters in length. Cyanobacteria in Pool 2 and Pool 3 display high levels of photosynthetic activity as revealed by high dissolved oxygen concentration (∼234 μM), low dissolved inorganic carbon concentrations, and the accumulation of visible O_2_ bubbles on the surface and within the fabric of the mat. Downstream pools (Pools 2 and 3) mix with seawater during high tide due to wave action, but this seawater influence does not appear to influence the Source Pool or Pool 1. Samples were collected and temperatures were measured at high tide, reflecting the lowest temperatures experienced by microbes in the pools—at low tide, hot spring input is dominant and temperatures rise (observed range at each site in Supplemental Table 1). Subaqueous surfaces in Pools 2 and 3 are covered in thick microbial mats. In Pool 2, the mat is coated in a layer of fluffy iron oxide similar to that in the Source Pool, with dense microbial mat below (Figure 2E). Pool 3 contains only patchy iron oxides, with mostly exposed microbial mats displaying a finger-like morphology. These “fingers” were 0.5-1 cm in diameter and up to ∼5 cm long and were closely packed and carpeting surfaces of Pool 3 below the high tide line, potentially related to turbulent mixing from wave action during high tide (Figure 2F). The Outflow is the outlet of a channel connecting Pool 2 to the bay. Its hydrology is dominantly marine with small admixtures of inflowing spring water (Figure 2G).

**Table 1:** Geochemistry of source water at Jinata Onsen.

|  |  |
| --- | --- |
| **T** | 63°C |
| **pH** | 5.4 |
| **DO** | 4.7 µM |
| **Fe<sup>2+</sup>** | 261 µM |
| **NH<sub>3</sub>/NH<sub>4</sub><sup>+</sup>** | 87 µM |
| **Cl<sup>-</sup>** | 654 mM |
| **SO<sub>4</sub><sup>2-</sup>** | 17 mM |
| **NO<sub>3</sub><sup>-</sup>** | <1.6 µM |
| **NO<sub>2</sub><sup>-</sup>** | <2.2 µM |
| **HPO<sub>4</sub><sup>-</sup>** | <1 µM |

**Figure 3:**
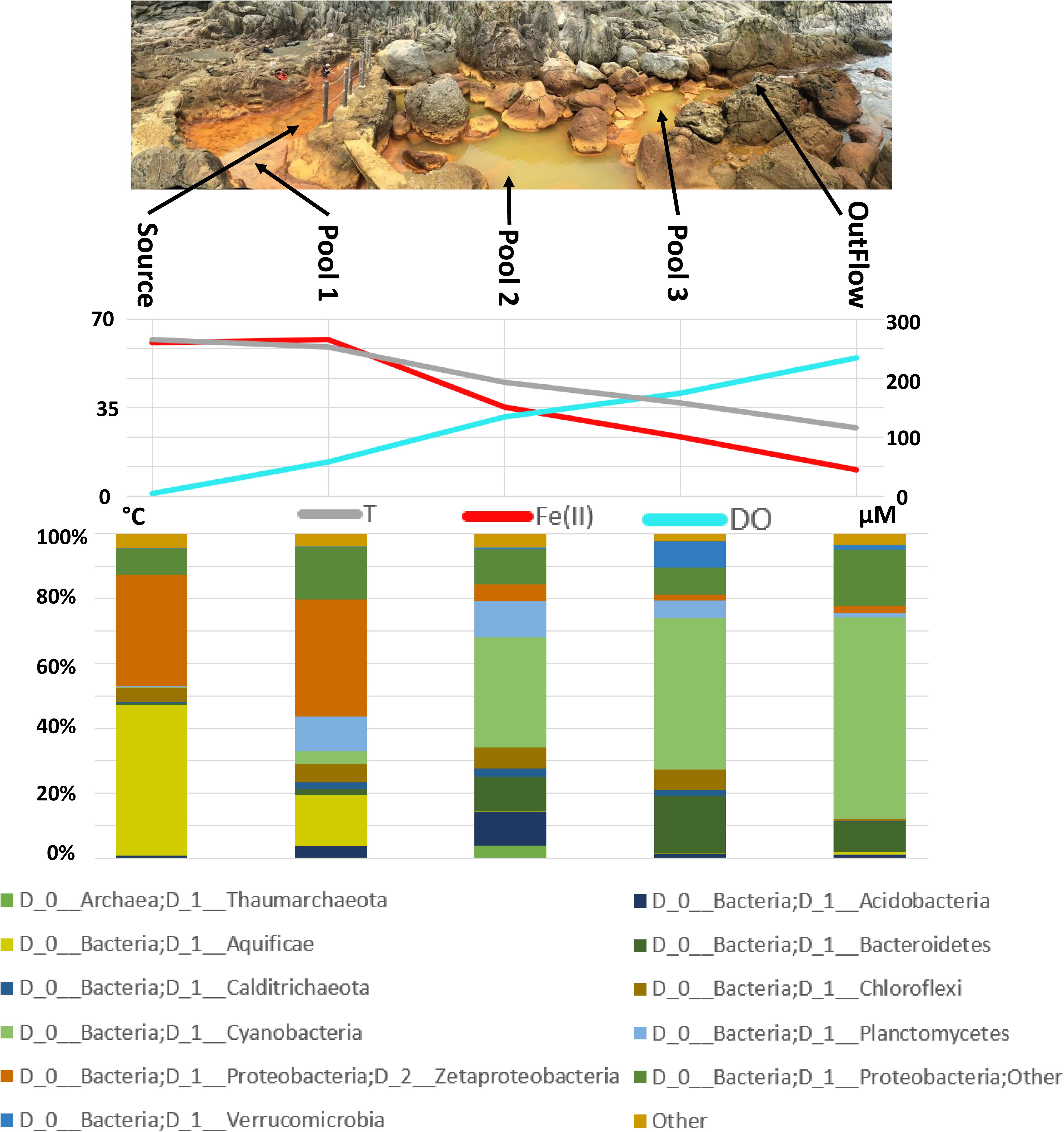
Summary of geochemical and microbiological trends along the flow path of Jinata Onsen. Top: panoramic view of Jinata Onsen, with Source Pool at left and flow of spring water toward the bay at right, with sampling locations indicated. Middle: geochemical transect across the spring, showing temperature (°C, left axis) and dissolved Fe(II) and O_2_ (μM, right axis). Bottom: stacked bar chart of relative community abundance of dominant microbial phyla as determined by 16S rRNA amplicon sequencing. Sequence data were binned at the phylum level and duplicate samples at each site were averaged. Reads that could not be assigned to a phylum were discarded; all phyla that do not make up more than 2% of the community at any one site have been collapsed to “Other”. Near the source, the community is predominantly made up of iron- and/or hydrogen-oxidizing organisms in the Proteobacteria and Aquificae phyla. As the hot spring water flows downstream, it equilibrates with the atmosphere and eventually mixes with seawater, resulting in downstream cooling, accumulation of oxygen, and loss of dissolved iron due to biological and abiotic processes. Oxygenic Cyanobacteria become progressively more abundant downstream Hydrogen- and iron-oxidizing lithotrophs dominate near the source, but phototrophic Cyanobacteria come to dominate downstream. Additional community diversity is found in Supplemental Table 4.

Jinata hot spring was visited twice for observation and community DNA sampling in 2016 (January and September), and again for observation and gas sampling in October 2017 and April 2018. These visits corresponded to a range of tidal conditions, including a spring low and high tide in September 2016. General features of the spring were consistent across this period (including abundance and distribution of iron minerals and microbial mats), differing primarily in an apparent tidal dependence in flow rate and water level of the spring and the amount of seawater influence on Pool 3. These differences in flow and mixing led to variation in water temperatures of 3-10 °C (Supplemental Table 1). At high tide, the flow rate of the spring increases, as does seawater influx to Pool 3. During the spring low tide, the spring flow stagnated and the water level of Source Pool and Pool 1 dropped by decimeters, with some portions draining entirely. During less extreme low tides observed on other dates, the spring flow was low but nonzero and the water level of the Source Pool did not drop significantly. While there is substantial variability in the flow rate from the spring based on tides (and resulting shifts in water level and temperature), the overall geochemistry of the source water and the microbial community appeared largely similar between expeditions.

### Geochemistry

Geochemical measurements along the flow path of Jinata Onsen revealed a major shift from hot, low-oxygen, high-iron source water to cooler, more oxygen-rich water with less dissolved iron downstream. Geochemistry measurements of Jinata source water are summarized in Table 1, while geochemical gradients along the stream outflow are summarized in Figure 3 and Supplemental Table 1. Source waters were slightly enriched in chloride relative to seawater (∼23.2 g/L in Jinata source water versus ∼19.4 g/L in typical seawater), depleted in sulfate (∼1.6 g/L in Jinata versus ∼2.7 g/L in seawater) but approached seawater concentrations downstream as mixing increased. Water emerging from the source was 63°C, very low in dissolved oxygen (∼4.7 μM), at pH 5.4, and contained substantial concentrations of dissolved iron (∼250 μM Fe^2+^). Dissolved organic carbon (DOC) in the source water was high (∼1.31 mM). It is unknown whether this is produced *in situ* or whether the source water emerges with high DOC. Both DOC and DIC decrease along the outflow of the spring (Supplemental Table 1). After emerging from the source, the spring water exchanges gases with the air due to mixing associated with water flow and gas ebullition, and DO rose to 39 μM at the surface of the Source Pool. As water flows downstream from the Source Pool, it cools slightly, exchanges gases with the atmosphere, and intermittently mixes with seawater below Pool 1.

Both H_2_ and CH_4_ were qualitatively detected in bubbles from the Source Pool following initial sampling in September 2016. However, during subsequent analyses to quantify the gas composition in October 2017 and April 2018 the gas was determined to contain CO_2_, CH_4_, N_2_ (Supplemental Table 2). This subsequent non-detection of H_2_ may be related to temporal variability in the gas composition at Jinata (e.g. following tidal influence; significant variability was observed in the CO_2_:N_2_ ratio between two sampling dates, Supplemental Table 2) or may reflect oxidation of H_2_ between sampling and analysis. The detection limit of H_2_ for these later measurements was ∼1 nmol/cc (in the gas phase of our quantitative gas analyses, or ∼1 nM in the aqueous phase, (3)), well above the energetic and ecological limits for hydrogenotrophic metabolisms (e.g. 53) leaving open the possibility of biologically significant H_2_ fluxes at Jinata around the time of sampling. The oxidation of H_2_ coupled to O_2_ reduction is a thermodynamically favorable process even at very low substrate concentrations (e.g. Δ_r_G’ < −375 kJ/mol with substrate concentrations of 0.1 nM H_2(aq)_ and 0.1 μM O_2(aq)_, well below our limit of detection) (32)). Consistent with this thermodynamic favorability, biology has been shown to make use of this metabolism in environments such as hot springs with H_2_ concentrations near our detection limits (21) and in Antarctic soils where microbes rely on uptake of trace atmospheric H_2_ at concentrations around 190 ppbv (53). Therefore the trace amounts of H_2_ which may be present in the source water at Jinata may be sufficient to support lithoautotrophy near the hot spring source in organisms possessing the genetic capacity for hydrogen oxidation as discussed below. Improved quantification of H_2_ concentrations and measurement of hydrogenase activity and the productivity of hydrogenotrophic microbes will be needed in future to determine the relative contribution of hydrogen oxidation to productivity at Jinata.

### 16S rRNA and genome-resolved metagenomic sequencing

16S rRNA and metagenomic sequencing of microbial communities at Jinata Onsen revealed a highly diverse community. In total, 16S rRNA amplicon sequencing recovered 456,737 sequences from the 10 samples at Jinata (Supplemental Tables 3-5). Reads per sample following filtering for quality and removal of chimeras ranged from 2,076 for Pool 3 Sample B to 96,268 for Pool 1 Sample A (median 32,222, mean 35,479, and standard deviation 26,014). On average 65% of the microbial community was recovered from Jinata samples at the 99% OTU level based on the Good’s Coverage statistic of the 16S rRNA gene (ranging from 50% coverage in the Outflow Sample A to 80% in the Pool 1 Sample A) and 82% at the 97% OTU level (69% for Pool 2 Sample B to 93% for the Pool 1 Sample B). MDS analysis (Supplemental Figure 1) demonstrates that samples from the same site are highly similar, and adjacent sites (e.g. Source Pool and Pool 1, Outflow and Pool 3) also show a high degree of similarity. However, there is a substantial transition in microbial community diversity between the most distant samples (e.g. Source Pool and Outflow).

Shotgun metagenomic sequencing of four samples from Jinata Onsen recovered 121 GB of data, forming a 1.48 Gb coassembly consisting of 1,531,443 contigs with an N50 of 1,494 bp. Nucleotide composition and differential coverage-based binning of the coassembly via multiple methods followed by dereplication and refinement resulted in a final set of 161 medium- or high-quality metagenome-assembled genomes (MAGs) following current standards (i.e. completeness >50% and contamination <10%) (7). These MAGs are from diverse phyla of Bacteria and Archaea (Figure 4); metagenome and MAG statistics with tentative taxonomic assignments for recovered MAGs are available in Supplementary Table 6, while MAGs of particular interest due to their potential contribution to primary productivity at this site or due to substantial genetic or metabolic novelty are discussed in depth below and shown in phylogenetic trees alongside reference strains in Figures 5-7.

**Figure 4:**
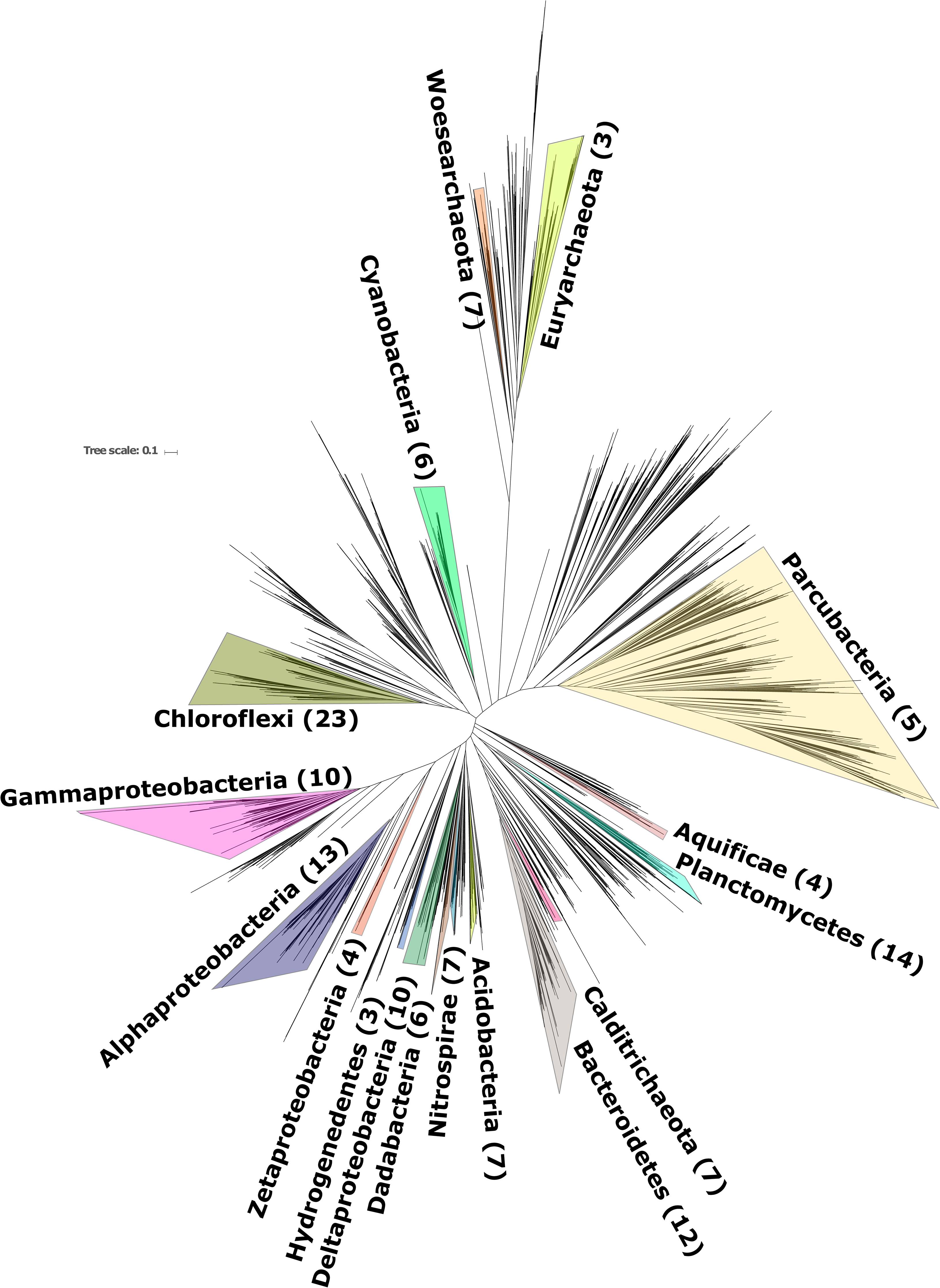
Phylogeny of Bacteria and Archaea based on concatenated ribosomal proteins. Numbers in parentheses next to phylum labels refer to number of MAGs recovered from Jinata Onsen. Labels for phyla with two or fewer MAGs recovered from Jinata omitted for clarity. The reference alignment was modified from Hug et al. (50). Full list of MAGs recovered available in Supplemental Table 6.

**Figure 5:**
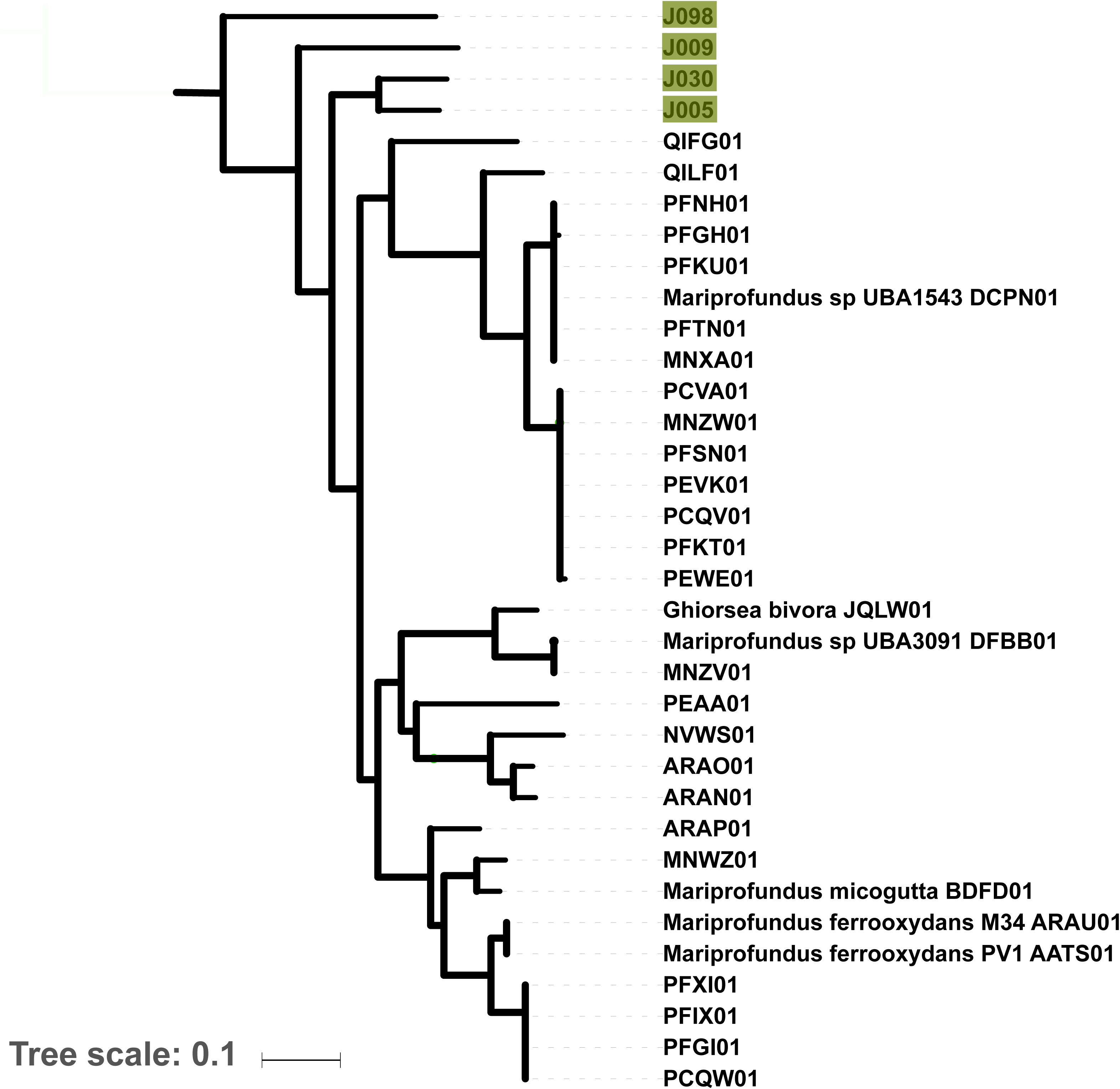
Phylogeny of the Zetaproteobacteria, rooted with Alphaproteobacteria, built with concatenated ribosomal protein sequences. Data from (80), (85), (105), and other draft genomes available on Genbank. All nodes recovered TBE support values greater than 0.7. In cases where reference genomes have a unique strain name or identifier, this is included; otherwise Genbank WGS genome prefixes are used.

## Discussion

As Jinata spring water flows from source to ocean, it transitions from hot, low-oxygen, high-iron water to cooler, iron-depleted, oxygen-rich water in downstream regions (Figure 3). Following this geochemical transition is a major shift in the composition of the microbial community, from a high-temperature, putatively lithotrophic community which produces little visible biomass upstream, to a lower temperature, community with well-developed, thick microbial mats downstream. This shift in community composition is summarized in Figure 3, with complete diversity data in the Supplemental Information (including OTU counts per samples in Supplemental Table 4 and relative abundance binned at the class level in Supplemental Table 5). Below, we discuss the overall physiological and taxonomic trends across the spring sites as inferred from diversity and genomic analysis.

### Potential for iron and hydrogen oxidation

The hot spring water emerging at the Source Pool at Jinata contains abundant dissolved Fe^2+^ and trace H_2_ (though measurements of gas content varied, as discussed above) (Table 1). Although rates of carbon fixation were not measured, the appearance of zetaproteobacterial veils and streamers and molecular evidence for lithoautotrophic microbes suggests that these electron donors may fuel productivity and determine the microbial community upstream at the Source Pool and Pool 1, where microbial mats are not well developed. The low accumulation of visible biomass in upstream regions of Jinata are similar to other microbial ecosystems fueled by iron oxidation (e.g. Oku-Okuhachikurou Onsen, 130, Fuschna Spring, 44, and Jackson Creek, 96), in which lithotrophic communities appear capable of accumulating less organic carbon than communities fueled by oxygenic photosynthesis (including those in downstream regions at Jinata).

Results of 16S rRNA sequencing indicate that the most abundant organisms in the Source Pool are members of the Aquificae family Hydrogenothermaceae (32% of reads in the Source Pool and 11.5% of reads in Pool 1). Members of this family of marine thermophilic lithotrophs are capable of iron and hydrogen oxidation, as well as heterotrophy (118) and may be utilizing Fe^2+^, H_2_, or dissolved organic carbon at Jinata. The seventh most abundant OTU in the Source Pool samples is a novel sequence 89% similar to a strain of *Persephonella* found in an alkaline hot spring in Papua New Guinea. *Persephonella* is a genus of thermophilic, microaerophilic hydrogen oxidizing bacteria within the Hydrogenothermaceae (38). Despite their abundance as assessed by 16S rRNA sequencing (Figure 3), only four partial Aquificae MAGs were recovered from Jinata of which only one (J026) was reasonably complete (∼94%). Two Aquificae MAGs recovered Group 1 NiFe hydrogenase genes, which may be used in hydrogenotrophy; the absence of hydrogenases from the other MAGs may be related to their low completeness, or could reflect a utilization of iron or other electron donors and not H_2_ in these organisms.

The other most abundant organisms near the source are members of the Zetaproteobacteria—a group typified by the neutrophilic, aerobic iron-oxidizing genus *Mariprofundus* common in marine systems (29). Zetaproteobacteria accounted for 24% of 16S rRNA sequences in the Source Pool and 26.5% in Pool 1. All Zetaproteobacteria characterized to date are obligate iron- and/or hydrogen-oxidizing lithoautotrophs (85), suggesting that these organisms may play a substantial role in driving carbon fixation in the Source Pool and Pool 1.

Members of the Mariprofundaceae have been observed to have an upper temperature limit for growth of 30 °C (30), while Zetaproteobacteria are found at Jinata at temperatures up to 63 °C. This currently represents a unique high-temperature environment for these organisms. In particular, the third most abundant OTU in the Source Pool and Pool 1 sample A is an unknown sequence that is 92% identical to a sequence from an uncultured zetaproteobacterium from a shallow hydrothermal vent in Papua New Guinea (82). This sequence likely marks a novel lineage of high-temperature iron-oxidizing Zetaproteobacteria.

The relative abundance of Hydrogenothermaceae drops off to less than 1% of sequences where microbial mats become well developed downstream of Pool 1, but Zetaproteobacteria continue to make up ∼1-4% percent of reads in Pool 2 and Pool 3 where dissolved iron concentrations are still significant (Figure 3). It may be that the relative abundance change is due more to the increase in abundance of other organisms, rather than a drop in the number of Zetaproteobacteria or their ability to make a living oxidizing iron. This hypothesis awaits confirmation by a technique such as qPCR. In contrast, the absence of Hydrogenothermaceae downstream may be a real signal driven by the rapid disappearance of trace H_2_ as an electron donor. However, in both cases, a drop in relative abundance is likely related to the increasing total biomass (i.e. number of cells) downstream as Cyanobacteria become more productive, leading to sequences from Hydrogenothermaceae and Zetaproteobacteria being diluted out by increased numbers of Cyanobacteria, Chloroflexi, and other sequences.

Four MAGs affiliated with the Zetaproteobacteria were recovered from Jinata with completeness estimates by CheckM ranging from ∼80 to ∼97% (J005, J009, J030, and J098). While these MAGs did not recover 16S rRNA genes, RpoB- and concatenated ribosomal protein-based phylogenies illustrated that members of this group at Jinata Onsen do not belong to the characterized genera *Mariprofundus* or *Ghiorsea*, but instead form separate basal lineages within the Zetaproteobacteria (Figure 5). Despite their phylogenetic distinctness, these MAGs largely recovered genes associated with aerobic iron oxidation as expected given the physiology of other Zetaproteobacteria. These include a terminal O_2_ reductase from the C-family of heme copper oxidoreductases for respiration at low O_2_ concentrations and Cyc2 cytochrome genes implicated in ferrous iron oxidation in Zetaproteobacteria and other taxa (e.g. Chlorobi) (41, 42, 61). Hydrogenase catalytic subunit genes (neither [NiFe] nor [FeFe]) were not recovered in zetaproteobacterial MAGs even at high completeness, suggesting that these organisms are not hydrogenotrophic. Consistent with the obligately autotrophic lifestyle of previously characterized Zetaproteobacteria, J009 and J098 encode carbon fixation via the Calvin cycle. However, J005 and J030 which did not recover genes for carbon fixation via the Calvin cycle (such as the large and small subunits of rubisco, phosphoribulose kinase, or carboxysome proteins). The high completeness of these MAGs (∼94-97%) makes it unlikely that these genes would all fail to be recovered (MetaPOAP False Negative estimates 10^-5^-10^-7^). The absence of carbon fixation pathways from these genomes together with the availability of abundant dissolved organic carbon in Pool 1 (∼1.3 mM) suggest that these organisms may be heterotrophic, a lifestyle not previously observed for members of the Zetaproteobacteria.

Seven MAGs were recovered from the enigmatic bacterial phylum Calditrichaeota (J004, J008, J042, J070, J075, J140, and J141) (Figure 6). While few members of Calditrichaeota have been isolated or sequenced, the best known of these is *Caldithrix abyssi* (84); this taxon was characterized as an anaerobic thermophile capable of lithoheterotrophic H_2_ oxidation coupled to denitrification and organoheterotrophic fermentation (1, 81). The Caldithrichaeota MAGs reported here are up to 97% complete (J004) and contain members with variable putative metabolic capabilities, potentially including aerobic hydrogen- or iron-oxidizing lithoautotrophy. In the Calditrichaeota MAGs recovered from Jinata Onsen, aerobic respiration via A-family heme copper oxidoreductases could potentially be coupled to autotrophic hydrogen oxidation (via the Group 1d NiFe hydrogenase in J042) or iron oxidation (via the *pioA* gene in J075); however, *Caldithrix abyssi* appears incapable of aerobic respiration despite encoding an A-family heme copper oxidoreductase (70). A MAG from a member of Calditrichaeota has previously been recovered from Chocolate Pots hot spring in Yellowstone National Park (34); together with the data presented here this suggests that this phylum may be a common member of microbial communities in iron-rich hot springs. Unlike previously described Calditrichaeota which are all heterotrophic (81), most of the Calditrichaeota MAGs reported here possess a putative capacity for carbon fixation via the Calvin cycle. J004 is closely related to *Caldithrix abyssi*, while the other MAGs form two distinct but related clades (Figure 6).

**Figure 6:**
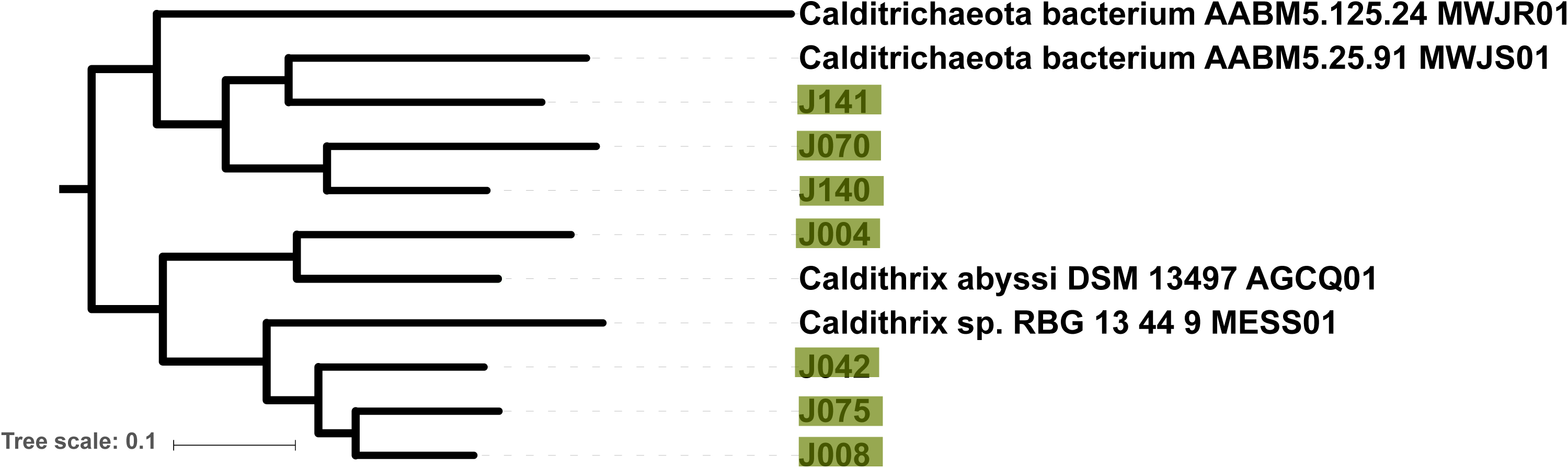
Phylogeny of the Calditrichaeota, rooted with Bacteroidetes, built with concatenated ribosomal protein sequences. Data from (70) and other draft genomes available on genomes have a unique strain name or identifier, this is included; otherwise Genbank WGS genome prefixes are used.

### Oxygenic photosynthesis

Cyanobacteria are nearly absent from near the Source Pool, but are observed in low numbers in Pool 1 and become abundant starting in Pool 2. The most abundant Cyanobacteria present are predominantly members of the Nostocales. This group includes *Leptolyngbya* and *Phormidium*, genera of filamentous non–heterocystous Cyanobacteria that are present in other hot springs of similar temperatures (e.g. 6, 98, 130). Diverse cyanobacterial MAGs were recovered, including members of the orders Pleurocapsales (J083), Chroococcales (J003 and J149), and Oscillatoriales (J007, J055, and J069). In the Outflow samples, chloroplast sequences are abundant, most closely related to the diatom *Melosira.*

Cyanobacteria are sometimes underrepresented in 16S rRNA amplicon sequencing datasets as a result of poor DNA yield or amplification biases (e.g. 88, 122), but the low abundance of Cyanobacteria near the Source Pool was confirmed by fluorescent microscopy, in which cells displaying cyanobacterial autofluorescence were observed abundantly in samples from the downstream samples but not in the Source Pool (Supplemental Figure 2). Thick microbial mats first appear in Pool 2 when Cyanobacteria become abundant, suggesting that oxygenic photosynthesis fuels more net carbon fixation than lithotrophy in these environments. Previously, it has been suggested that high ferrous iron concentrations are toxic to Cyanobacteria, and that this would have greatly reduced their productivity under ferruginous ocean conditions such as those that may have persisted through much of the Archean era (117). The abundant Cyanobacteria observed to be active at Jinata under high iron concentrations suggest that Cyanobacteria can adapt to ferruginous conditions, and therefore iron toxicity might not inhibit Cyanobacteria over geological timescales. Indeed, the soluble iron concentrations observed at Jinata are higher (150-250 μM) than predicted for the Archean oceans (<120 μM, 49) or observed at other iron-rich hot springs (∼100-200 μM, 91, 130), making Jinata an excellent test case for determining the ability of Cyanobacteria to adapt to high iron concentrations. Culture-based physiological experiments may be useful to determine whether Jinata Cyanobacteria utilize similar strategies to other iron-tolerant strains (e.g. by those in Chocolate Pots Hot Spring, 91, or the ferric iron-tolerant *Leptolyngbya*-relative *Marsacia ferruginose*, 9) or whether Jinata strains possess unique adaptations that allow them to grow at higher iron concentrations than known for other environmental Cyanobacteria strains. This will in turn provide insight into whether iron tolerance is due to evolutionarily conserved strategies or whether this is a trait that has evolved convergently multiple times.

### Diverse novel Chloroflexi from Jinata Onsen

In addition to the primary phototrophic and lithotrophic carbon fixers at Jinata, 16S rRNA and metagenomic data sets revealed diverse novel lineages within the Chloroflexi phylum. A total of 23 Chloroflexi MAGs were recovered, introducing substantial genetic and metabolic diversity that expands our understanding of this group. While the best known members of this phylum are Type 2 Reaction Center-containing lineages such as *Chloroflexus* and *Roseiflexus* within the class Chloroflexia (e.g. 121), phototrophy is not a synapomorphy of the Chloroflexi phylum or even the Chloroflexia class (e.g. 126) and most of the diversity of the phylum belongs to several other classes made up primarily of nonphototrophic lineages (131). The bulk of Chloroflexi diversity recovered from Jinata belongs to “subphlyum I”, a broad group of predominantly nonphototrophic lineages that was originally described based on the class- or order-level lineages Anaerolineae and Caldilineae (141), but also encompasses the related groups Ardenticatenia, Thermoflexia, and *Candidatus* Thermofonsia (22, 63, 131).

16S rRNA analysis indicates that members of Anaerolineae and *Candidatus* Thermofonsia (annotated by Silva and GTDB-Tk as the order SBR1031) are fairly abundant at Jinata, with Anaerolineae at ∼3% relative abundance in the Source and Pool 1 and *Ca.* Thermofonsia at ∼3.5% relative abundance in Pool 2 and Pool 3. Three MAGs recovered from Jinata (J082, J097, and J130) are associated with the Anaerolineae class as determined by RpoB and concatenated ribosomal protein phylogenies, along with seven associated with *Ca.* Thermofonsia (J027, J033, J036, J038, J039, J064, and J076). Particularly notable among these MAGs is J036, a close relative of the phototrophic *Ca.* Roseilinea gracile (68, 119, 120). J036 contains a 16S rRNA gene that is 96% similar to that of *Ca*. Roseilinea gracile, and two-way AAI estimates (97) showed 73.6% similarity between the two strains, indicating these strains are probably best classified as distinct species within the same genus. Unlike other phototrophs in the Chloroflexi phylum that are capable of photoautotrophy via the 3-hydroxypropionate bicycle or the Calvin Cycle (67, 102), J036 and *Ca.* Roseilinea gracile do not encode carbon fixation and are likely photoheterotrophic. Previous analyses suggested that the Roseilinea lineage belongs to the Anaerolineae (68) or Thermofonsia (131); however, our updated phylogeny presented here places J036 and Roseilinea in a separate lineage along with J033 and J162, diverging just outside of the Anaerolineae+Thermofonsia clade, suggesting that these strains may instead be yet another class- or order-level lineage within the broader “subphylum I” of Chloroflexi (Figure 7), an interpretation supported by analysis via GTDB-Tk which places these genomes outside of characterized clades (Supplemental Table 6).

**Figure 7:**
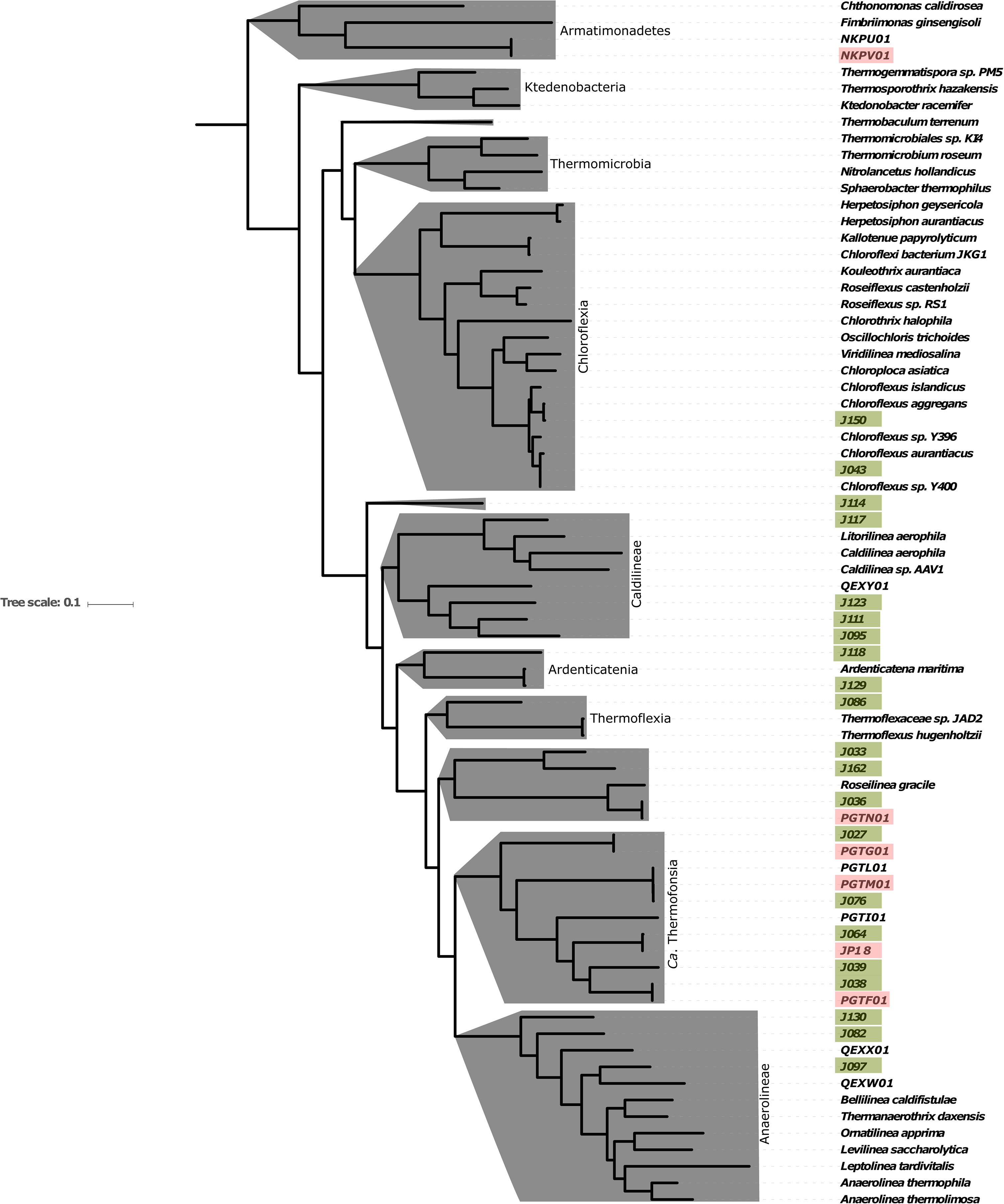
Detailed phylogeny of the Chloroflexi phylum, with class-level clades highlighted in gray, built with concatenated ribosomal protein sequences. The large basal class Dehalococcoidia, which was not observed in 16S rRNA or metagenome data from Jinata, is omitted for clarity. The phylogeny contains MAGs reported here, members of the Chloroflexi phylum previously described (18, 22, 40, 43, 45, 46, 47, 64, 72, 87, 108, 126, 127, 131, 132, 133), and members of the closely related phylum Armatimonadetes as an outgroup (23, 129). MAGs described here are highlighted in green, MAGs previously reported from Jinata Onsen highlighted in pink. All nodes recovered TBE support values greater than 0.7. In cases where reference genomes have a unique strain name or identifier, this is included; otherwise Genbank WGS genome prefixes are used.

The Chloroflexi class Ardenticatenia was first described from an isolate from an iron-rich Japanese hydrothermal field (63) and has since been recovered from sulfidic hot springs as well (132). Members of Ardenticatenia were present at up to 1.2% relative abundance in Pool 3 in 16S amplicon data. A MAG closely related to *Ardenticatena maritima* was recovered from Jinata Onsen, J129. While *Ardenticatena maritima 110S* contains a complete denitrification pathway (47), MAG J129 did not recover any denitrification genes. This could be related to the relatively low completeness of this MAG (∼70%), but False Negative estimates by MetaPOAP (134) indicates that the probability that all four steps in the canonical denitrication pathway would fail to be recovered in J129 given their presence in the source genome is less than 0.8%, suggesting that most if not all denitrification genes are absent and that the capacity for denitrification is not universal within members of *Ardenticatena.* This would be consistent with broad trends in the apparently frequent modular horizontal gene transfer of partial denitrification pathways between disparate microbial lineages to drive rapid adaption and metabolic flexibility of aerobic organisms in microoxic and anoxic environments, for reasons that are still not well established (19, 113).

Members of the Chloroflexi class Caldilineae were present at up to 0.5% abundance at Jinata in the 16S rRNA dataset. Members of the Caldilineae have previously been isolated from intertidal hot springs in Iceland (57) and Japanese hot springs (99). Characterized organisms in this class are filamentous, anaerobic, or facultatively aerobic heterotrophs (39, 57, 99); and therefore these taxa may play a role in degrading biomass within low-oxygen regions of microbial mats at Jinata. Three MAGs were recovered that form a deeply branching lineage with the Caldilineae class (J095, J111, and J123), sister to the previously characterized genera *Caldilinea* and *Litorilinea.* Like other members of the Caldilineae, these strains encode aerobic respiration via A-family heme copper oxidoreductases and both a *bc* complex III and an alternative complex III, and are therefore likely at least facultatively aerobic. J095 also encodes carbon fixation via the Calvin cycle as well as a Group 1f NiFe hydrogenase, suggesting a potential capability for lithoautotrophy by hydrogen oxidation, expanding the known metabolic diversity of this class and the Chloroflexi phylum as a whole.

MAG J114 branches at the base of subphylum I of the Chloroflexi, potentially the first member of a novel class-level lineage. The divergence between Anaerolineae and Caldilineae has been estimated to have occurred on the order of 1.7 billion years ago (102). The phylogenetic placement of J114 suggests that it diverged from other members of subphylum I even earlier, and it may be a good target for future investigation to assess aspects of the early evolution of the Chloroflexi phylum. J114 encodes aerobic respiration via an A-family heme copper oxidoreductase and an alternative complex III like many other nonphototrophic Chloroflexi lineages (e.g. 126, 131) as well as a Group 1f NiFe hydrogenase and carbon fixation via the Calvin Cycle, suggesting the capacity for aerobic hydrogen-oxidizing autotrophy—a lifestyle not previously described for members of the Chloroflexi.

### Conclusions

To our knowledge, this is the first overall geomicrobiological characterization of Jinata Onsen, providing baseline descriptions of geochemistry and microbial diversity in order to establish a series of testable hypotheses which can be addressed by future studies. We have also provided genome-resolved metagenomics sequencing of this site focusing on members of the microbial community predicted to be responsible for the bulk of primary productivity in this system along with other organisms belonging to novel or under-characterized lineages. However, this is just a subset of the diverse microbial populations at Jinata Onsen; many more MAGs from across the tree of life were recovered than are discussed in detail here but which may be of use to others (Figure 4, Supplemental Table 6).

The diversity of iron oxidizing bacteria at Jinata is different than in other Fe^2+^-rich springs and environments. For example, in freshwater systems such as Oku-Okuhachikurou Onsen in Akita Prefecture, Japan (130), and Budo Pond in Hiroshima, Japan (60), iron oxidation is driven primarily by the activity of chemoautotrophs such as members of the Gallionellaceae. In contrast, at Chocolate Pots hot spring in Yellowstone National Park, USA, iron oxidation is primarily abiotic, driven by O_2_ produced by Cyanobacteria, with only a small contribution from iron oxidizing bacteria (34, 123). The distinct iron-oxidizing community at Jinata Onsen may be related to the salinity of the spring water, or biogeographically by access to the ocean, as Zetaproteobacteria are typically found in marine settings, particularly in deep ocean basins associated with hydrothermal iron sources (30). Despite the taxonomically distinct iron oxidizer communities between Jinata and Oku-Okuhachikurou Onsen, both communities support only limited visible biomass in regions dominated by iron oxidizers (130), perhaps reflecting the shared biochemical and bioenergetic challenges of iron oxidation incurred by diverse iron oxidizing bacteria including Gallionellaceae and Zetaproteobacteria (5, 30, 130). Future work focused on isolation and physiological characterization of microbes, quantification of rates and determination of microbial drivers of carbon fixation and aerobic and anaerobic heterotrophy, and carbon isotope profiling of organic and inorganic species along the flow path of the hot spring will be necessary to fully characterize the activity of microbes at Jinata and to fully compare this system to other areas with high dissolved ferrous iron concentrations (e.g. Oku-Okuhachikurou Onsen, 130, Fuschna Spring, 44, Jackson Creek, 96, and Chocolate Pots Hot Spring, 34, 123).

The relatively high concentrations of dissolved organic carbon (DOC) measured in Pool 1 (∼1.3 mM) may stimulate heterotrophic activity by the microbial community at Jinata, coupled to aerobic or anaerobic respiration (such as dissimilatory iron reduction, as observed in other iron-rich hot springs, e.g. 33), resulting in the drawdown of DOC downstream. The source of this DOC is unclear; future work will be necessary to determine whether DOC is present in the source water or if it is produced *in situ* by the microbial community in the Source Pool and Pool 1. Future work is also needed to evaluate the potential for dissimilatory iron reduction and other anaerobic metabolisms at this site.

Throughout Earth history, the metabolic opportunities available to life, and the resulting organisms and metabolisms responsible for driving primary productivity, have been shaped by the geochemical conditions of the atmosphere and oceans. The modern, sulfate-rich, well-oxygenated oceans we see today reflect a relatively recent state—one typical of only the last few hundred million years (e.g. 79). For the first half of Earth history, until ∼2.3 billion years ago (Ga), the atmosphere and oceans were anoxic (54), and the oceans were largely rich in dissolved iron but poor in sulfur (124). At this time, productivity was low and fueled by metabolisms such as methanogenesis and anoxygenic photosynthesis (14, 66, 135). Following the expansion of oxygenic photosynthesis by Cyanobacteria and higher primary productivity around the Great Oxygenation Event ∼2.3 Ga (20, 31, 128, 137), the atmosphere and surface ocean accumulated some oxygen, and the ocean transitioned into a state with oxygenated surface waters but often anoxic deeper waters, rich in either dissolved iron or sulfide (13, 55, 56, 92). At Jinata Onsen, this range of geochemical conditions is recapitulated over just a few meters, providing a useful test case for probing the shifts of microbial productivity over the course of Earth history. In particular, the concomitant increase in visible biomass at Jinata as the community shifts from lithotrophy toward water-oxidizing phototrophy (i.e. oxygenic photosynthesis) is consistent with estimates for greatly increased primary production following the evolution and expansion of Cyanobacteria around the GOE (20, 101, 106, 128, 135, 137).

The dynamic abundances of redox-active compounds including oxygen, iron, and hydrogen at Jinata may not only be analogous to conditions on the early Earth, but may have relevance for potentially habitable environments on Mars as well. Early Mars is thought to have supported environments with metabolic opportunities provided by the redox gradient between the oxidizing atmosphere and abundant electron donors such as ferrous iron and molecular hydrogen sourced from water/rock interactions (e.g. 51), and production of these substrates may continue today (24, 111), potentially supporting past or present life in the Martian subsurface (112). Understanding the potential productivity of microbial communities fueled by lithotrophic metabolisms is critical for setting expectations of the presence and size of potential biospheres on other worlds and early in Earth history (e.g. 135, 136, 137). Uncovering the range of microbial metabolisms present under the environmental conditions at Jinata, and their relative contributions to primary productivity, may therefore find application to predicting environments on Mars most able to support productive microbial communities.

## Data availability

Raw 16S rRNA, raw metagenomic sequence data, and MAGs have been uploaded and made publicly available on NCBI under Project Number PRJNA392119 (genome accession numbers can be found in Supplemental Table 6).

## Acknoweldgements

LMW acknowledges support from NASA NESSF (#NNX16AP39H), NSF (#OISE 1639454), NSF GROW (#DGE 1144469), Lewis and Clark Fund for Exploration and Field Research in Astrobiology, the ELSI Origins Network, and an Agouron Institute postdoctoral fellowship. WWF acknowledges the support of NASA Exobiology award #NNX16AJ57G, the David and Lucile Packard Foundation, and the Simons Collaboration on the Origins of Life. MN and YU acknowledge the support of the Japan Society for the Promotion of Science (JP17K14412, JP17H06105)

---

**Supplemental Figure 1:**
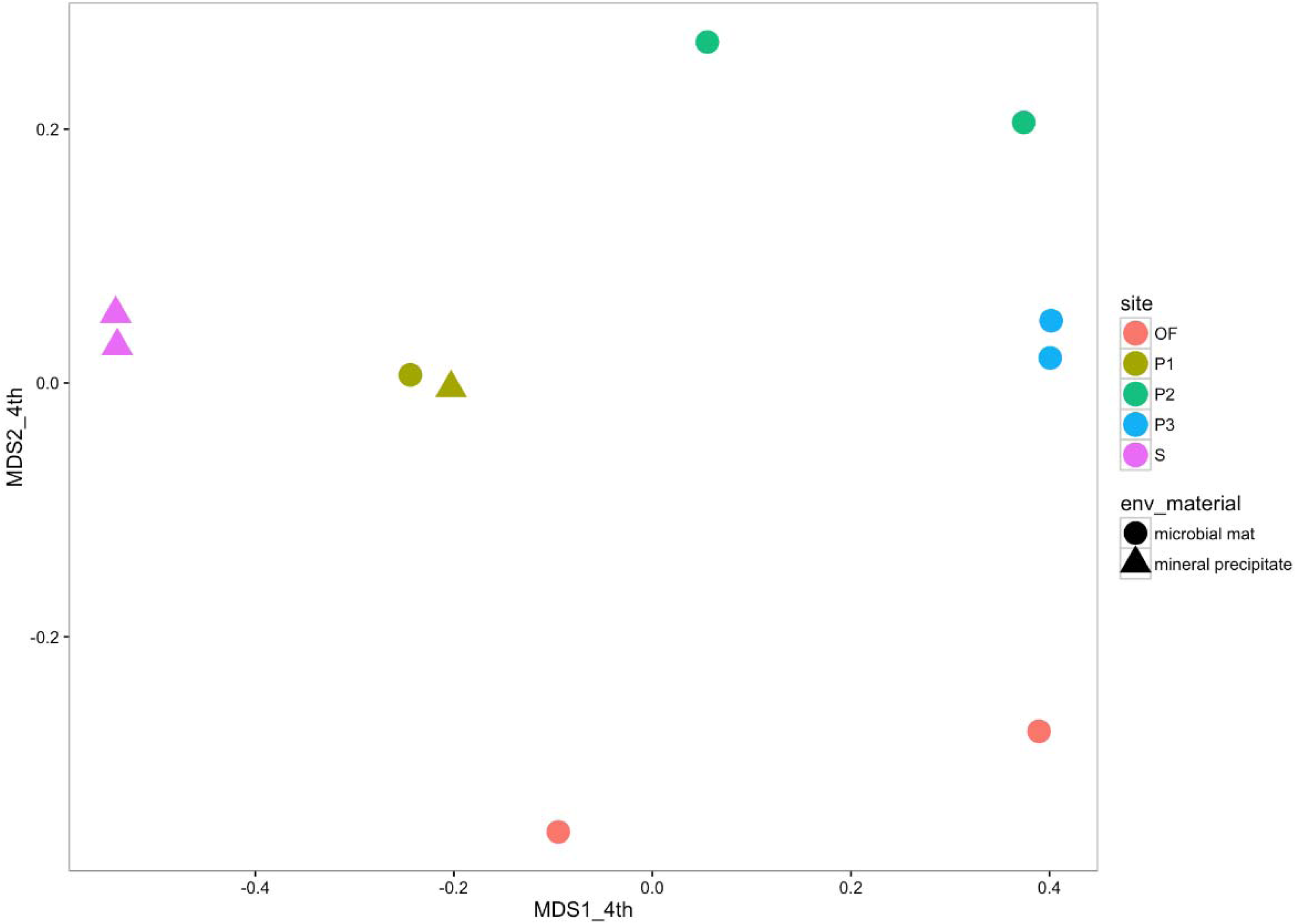
Multidimensional scaling plot of Jinata samples. Each point represents the recovered microbial community from a given sample, with sites identified by color and sample type by shape. Samples plotting close to each other are relatively more similar in community composition. Abundance data are transformed by the 4^th^ root to down-weight the effect of abundant taxa. Stress value is 0.0658.

**Supplemental Figure 2:** Microscopy images of sediment (Source Pool and Pool 1) or mat (Pool 2, Pool 3, and Out Flow). Left are light microscopy images. Center and right are fluorescence images. At center, blue signal is DAPI-stained (Excitation: 365nm, Emission: BP445∼50nm). At right, red is autofluorescence signal of Cyanobacteria (BP395∼440nm, LP470nm). Scale bars 50 μm.

**Supplemental Table 1:** Geochemistry and brief description at sampling sites along the flow path of Jinata Onsen as discussed in the text.

| | pH | T<br>(°C) | Fe(II)<br>(μM) | DO<br>(μM) | DIC (mM) | DOC (mM) | Descriptions |
| --- | --- | --- | --- | --- | --- | --- | --- |
| **Source Pool** | **5.4** | **60-63** | **260** | **4.7 (source) 39 (surface)** | **Not measured** | **Not measured** | **Fluffy red iron oxide precipitate** |
| **Pool 1** | **5.8** | **59-59.5** | **265** | **58** | **5.51 ± 0.28** | **1.31 ± 0.18** | **Reddish precipitate and streamers in shallower regions, more yellowish deeper** |
| **Pool 2** | **6.5** | **44.5-54** | **151** | **134** | **2.09 ± 0.11** | **0.76 ± 0.10** | **Iron oxide-coated microbial mats. Orange to orange-green.** |
| **Pool 3** | **6.7** | **37.3-46** | **100** | **175** | **1.79 ± 0.09** | **0.70 ± 0.10** | **Green or mottled orange-green microbial mats, commonly with 1-5cm finger-like morphology.** |
| **Outflow** | **6.5** | **27-32** | **45** | **234** | **Not measured** | **Not measured** | **Ocean water within mixing zone at high tide, with constant flow of spring water from Pool 2. Thin green microbial mats.** |

**Supplemental Table 2:**
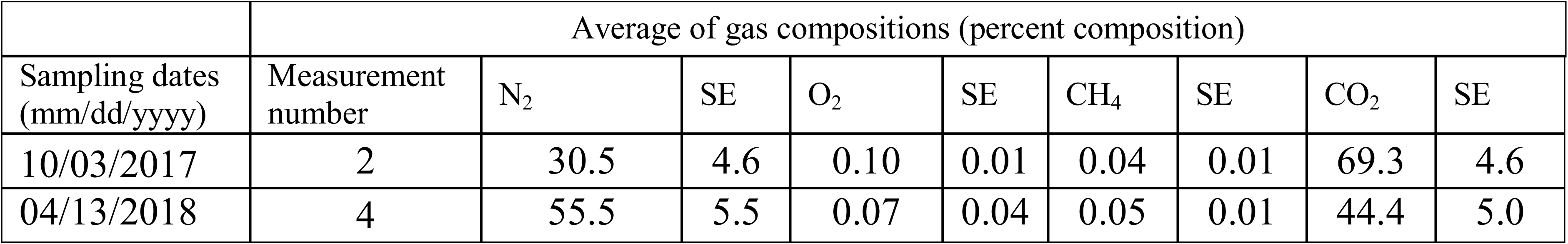
Gas composition of bubbles collected from the Source Pool at Jinata Onsen.

**Supplemental Table 3:** Diversity metrics of Jinata sequencing. Diversity metrics calculated for both 99% and 97% sequence identity cutoffs for assigning OTUs.

| Sample: | Reads: | OTUs<br>(99%): | Good<br>Coverage<br>(99%): | Shannon<br>Index<br>(99%): | Inverse<br>Simpson<br>(99%): | OTUs<br>(97%): | Good<br>Coverage<br>(97%): | Shannon<br>Index<br>(97%): | Inverse<br>Simpson<br>(97%): |
| --- | --- | --- | --- | --- | --- | --- | --- | --- | --- |
| **Source**<br>**Pool A** | 48680 | 18832 | 0.665 | 10.443 | 35.146 | 6951 | 0.907 | 7.996 | 17.361 |
| **Source**<br>**Pool B** | 18235 | 7772 | 0.646 | 10.388 | 68.018 | 4139 | 0.844 | 8.651 | 31.822 |
| **Pool 1 A** | 96268 | 25305 | 0.788 | 10.172 | 53.546 | 11734 | 0.920 | 8.403 | 28.691 |
| **Pool 1 B** | 56672 | 13835 | 0.797 | 8.813 | 26.975 | 5598 | 0.933 | 6.818 | 13.827 |
| **Pool 2 A** | 35690 | 12489 | 0.713 | 9.625 | 22.248 | 7352 | 0.855 | 8.271 | 16.600 |
| **Pool 2 B** | 4454 | 2274 | 0.560 | 9.066 | 49.599 | 1729 | 0.689 | 8.104 | 27.390 |
| **Pool 3 A** | 28273 | 11334 | 0.665 | 10.046 | 35.705 | 6766 | 0.824 | 8.403 | 20.034 |
| **Pool 3 B** | 2076 | 1166 | 0.522 | 8.832 | 75.900 | 786 | 0.699 | 7.489 | 35.312 |
| **Outflow**<br>**A** | 31980 | 18486 | 0.497 | 11.989 | 64.318 | 11994 | 0.712 | 10.538 | 34.881 |
| **Outflow**<br>**B** | 32465 | 10896 | 0.713 | 9.281 | 25.691 | 5918 | 0.857 | 7.133 | 11.585 |

**Supplemental Table 4:** 16S rRNA data as OTU table with sequences.

**Supplemental Table 5:** 16S rRNA data as relative abundance binned at the class level.

**Supplemental Table 6:** High- and medium-quality metagenome-assembled genomes (MAGs) (>50% completeness and <10% contamination) recovered from Jinata Onsen. Predicted taxonomy based on placement in reference phylogeny as presented in Figure 4 and by GTDB-Tk (90). Optimal growth temperatures predicted following methods from (78).

**Supplemental Table 7:** Presence of genes involved in aerobic respiration, hydrogen- and iron-oxidation, and carbon fixation in MAGs discussed in the text.

