## Supplementary figures and images for "Geochemical and metagenomic characterization of Jinata Onsen, a Proterozoic-analog hot spring, reveals novel microbial diversity including iron-tolerant phototrophs and thermophilic lithotrophs"

### SupplementalFigure2

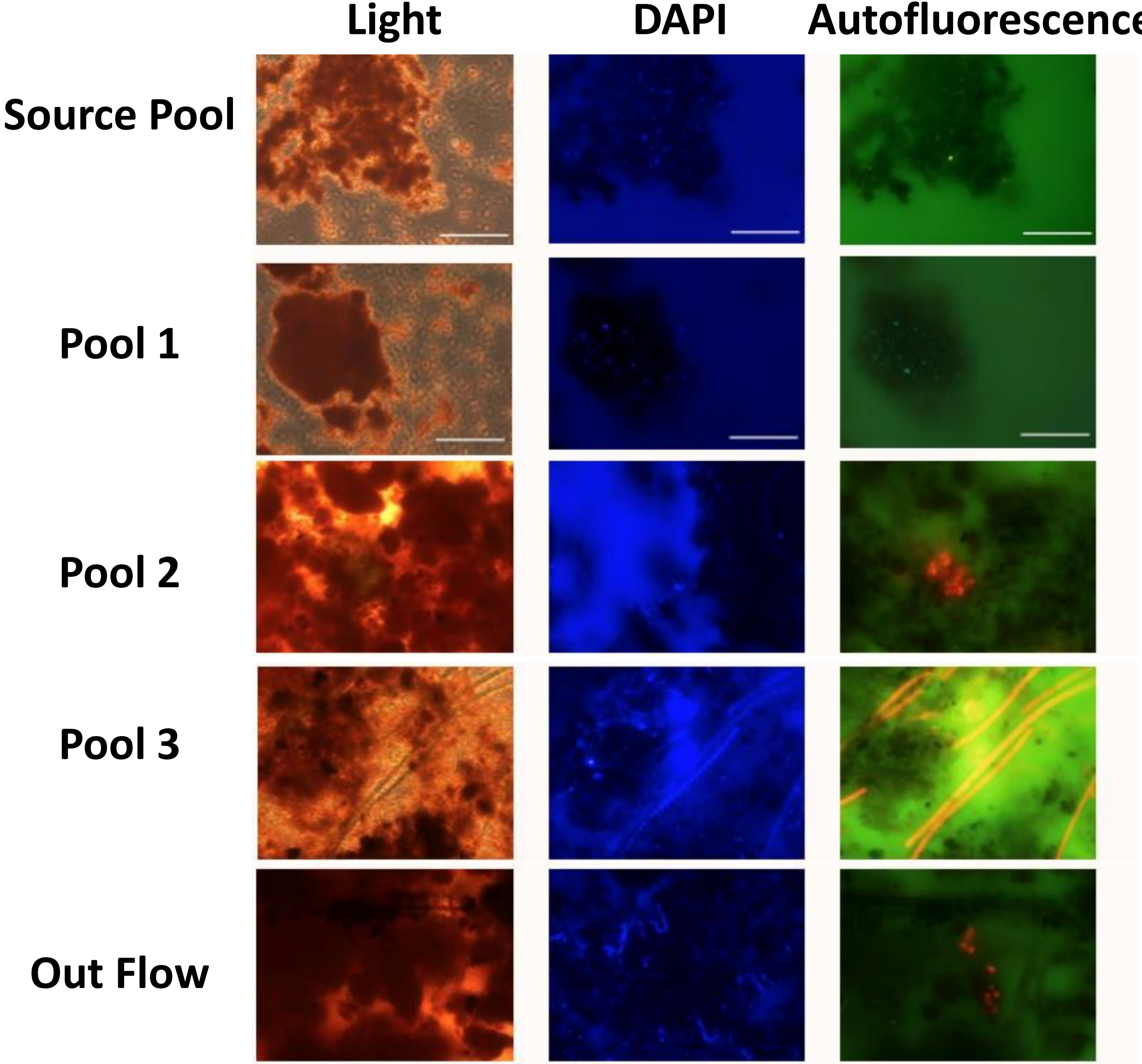
